# Immune-Epithelial Interactions via TGF-β Orchestrates Stem-Cell Niche Formation and Morphogenesis

**DOI:** 10.1101/2025.05.22.655596

**Authors:** Nicole Traugh, Colin Trepicchio, Gat Rauner, Meadow Parrish, Charlotte Kuperwasser

## Abstract

Mesenchymal-epithelial interactions are well-established for organogenesis, but the role of immune cells in these processes remains less explored. Organoids offer powerful models for studying tissue development, especially where in vivo models are limited. In this study, we established an immune-epithelial co-culture model using an advanced 3D hydrogel-based breast organoid model that enables real-time investigation of human breast organogenesis. We employed live imaging and cell tracking techniques to visualize the induction process and the formation of an immune-mediated niche. Incorporation of immune cells into the organoid structure demonstrates their dual role as inducers of epithelial proliferation and as a persistent component of the stem cell niche. In addition, co-culturing epithelial cells with tissue-resident immune cells from healthy donors, particularly those enriched in double negative T cells, significantly accelerates organoid development and increases organoid formation through TGF-β signaling. These findings highlight a previously unappreciated role for immune-epithelial crosstalk in early tissue development and provide insights into the dynamic nature of stem cell niches.

## Introduction

The formation of complex tissues during development depends on tightly regulated cell-cell interactions and signaling pathways. While mesenchymal-epithelial interactions are well-recognized as fundamental drivers of organogenesis, the potential role of immune cells in these developmental processes has been comparatively underexplored. Traditionally viewed through the lens of immune surveillance and pathogen defense, immune cells are increasingly being recognized for their contributions to tissue homeostasis, repair, and regeneration^1–7^.

Organoids, three-dimensional structures derived from stem or progenitor cells that recapitulate key features of native tissue architecture and function, have emerged as powerful models for studying human development and disease. These models allow for controlled investigation of cellular dynamics in a physiologically relevant context, providing insights into processes that are difficult to study in vivo. To date, most organoid systems have focused on epithelial and mesenchymal components, often overlooking the potential contributions of immune cells.

Immune cells, including various subsets of T cells, play critical roles in modulating epithelial behavior during tissue repair and homeostasis^8–10^. Among these, γδ T cells are known for their rapid response to stress signals and their ability to produce cytokines and growth factors that influence epithelial proliferation and differentiation. During development, γδ T cells are among the first cells to develop in the embryonic thymus, often appearing before conventional αβ T cells^11^. In humans, γδ T cells can be detected in the by 8-9 weeks of gestation where they migrate to peripheral tissues such as the skin, gut, and lungs, by 13-16 weeks of gestation^12,13^. Their early migration to epithelial sites suggests a role in tissue development and remodeling, yet it is unclear if and how they may play a role in organ formation.

In this study, we investigate the role of immune cells during breast organoid development. By incorporating tissue-resident immune cells into an advanced 3D hydrogel-based organoid model, we were able to observe immune-epithelial interactions in real time. Our findings reveal that immune cells from donor samples enriched in γδ T cells, significantly accelerate organoid formation and maturation through TGF-β signaling. Furthermore, the incorporation of immune cells into the organoid structure suggests that they function not only as inducers of epithelial proliferation but also as persistent components of the stem cell niche.

These observations provide new insights into the dynamic interplay between immune cells and epithelial tissues during development and highlight the broader implications of immune-epithelial crosstalk in tissue homeostasis, regenerative medicine, and disease progression.

## Results

### A 3D Organoid Hydrogel Co-Culture to Study Epithelial-Immune Cell Interactions

Co-culturing epithelial stem cells and immune cells together in a 3D culture system presents challenges due to differing growth requirements, the need for a suitable extracellular matrix (ECM) for both cell types, and the complexity of monitoring cell-cell interactions. Therefore, the first step was to evaluate whether the hydrogel system used to culture breast organoids could support such a co-culture. The 3D hydrogel scaffold is designed to induce niche signals and promote differentiation of single breast stem cells within a controlled ECM environment^14–17^. We aimed to assess whether it could also support the incorporation of tissue-resident immune cells to establish a co-culture system that might facilitate detailed observation of epithelial and immune cell interactions.

Immune and epithelial cells were co-isolated from disease-free human breast tissue reduction mammoplasty (N=12 donors). Each tissue sample was enzymatically dissociated into single-cell suspension, after which CD45^+^ immune cells were selectively separated from the remaining epithelial and stromal populations using releasable magnetic beads conjugated to CD45 antibodies (Fig. 1a). Following magnetic bead removal, the isolated CD45^+^ immune cells and matched breast epithelial cells from the same donor were collected and embedded into a hydrogel ECM composed of Collagen 1, hyaluronic acid, fibronectin and laminin. Co-cultured hydrogels were grown in serum-free media minimally supplemented with hEGF, hydrocortisone, insulin, and bovine pituitary extract (Fig 1a). To visualize and monitor immune cells in 3D, CD45+ cells were labeled with Cytopainter green dye that is retained only in viable cells and becomes progressively diluted as cells divide. Twenty-one days post seeding, bright green fluorescence remained, indicating that CD45^+^ immune cells survived within the hydrogel but underwent little to no proliferation. (Fig 1b).

**Figure 1:**
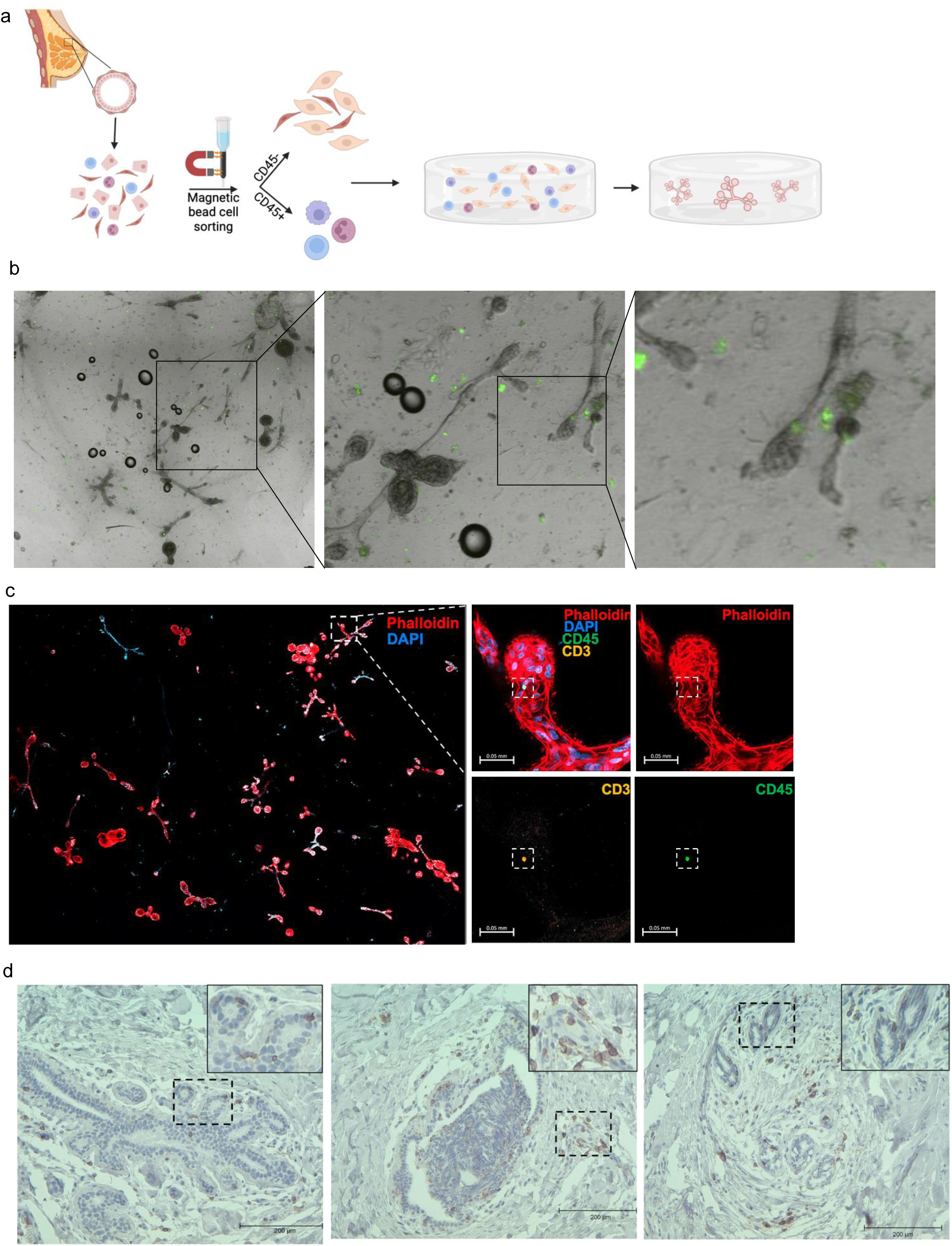
Establishment of patient-matched breast epithelial and immune cell 3D hydrogel co-culture system. (a) Schematic overview of the workflow for isolation of primary human breast epithelial and immune cells from reduction mammoplasty patients and embedding into 3D hydrogels for co-culture. (b) Representative brightfield images showing persistence of green CD45+ immune cells in hydrogels after 21 days of culture. (c) Immunofluorescent (IF) images of co-cultured 3D hydrogels after 21 days of culture, stained with phalloidin (red) for actin cytoskeleton, DAPI (blue) for nuclei, CD45 (green) for immune cells, and CD3 (orange) for T cells. Scale bar = 500µm. (d) Immunohistochemistry (IHC) images of human breast tissue (n=3). Images show immune cells localize both within epithelial ducts and in the surrounding stromal regions in vivo. Scale bar = 200µm.

We next assessed how the two populations spatially organized in co-culture. By 21-28 days epithelial cells had formed terminal ductal lobule units (TDLUs). Throughout this period, CD45^+^ immune cells stayed in close proximity to, and were often incorporated into, the developing organoids (Fig 1c, Supplemental Fig. 1a-c). This suggests an active, long-term role for immune cells in morphogenesis, rather than mere passive association. This intralobular localization mirrors that seen in vivo within human breast tissue (Fig 1d), underscoring that the hydrogel ECM simulates a natural environment conducive to both epithelial and immune cell interactions, providing a valuable context for studying immune cell behavior within breast tissue.

### Immune Cells Stimulate Organoid Formation

Building on the observation that CD45^+^ immune cells survive, remain proximal to, and ultimately integrate into developing TDLUs within the hydrogel matrix (Fig. 1b-e), we next asked whether this intimate association confers functional benefits to epithelial morphogenesis. Using the donor-matched co-culture system, we seeded immune and epithelial cells at a ratio of 1:10 and quantified three readouts of morphogenesis over time: total organoid number, duct number, and epithelial area (Figure 2a-d).

**Figure 2:**
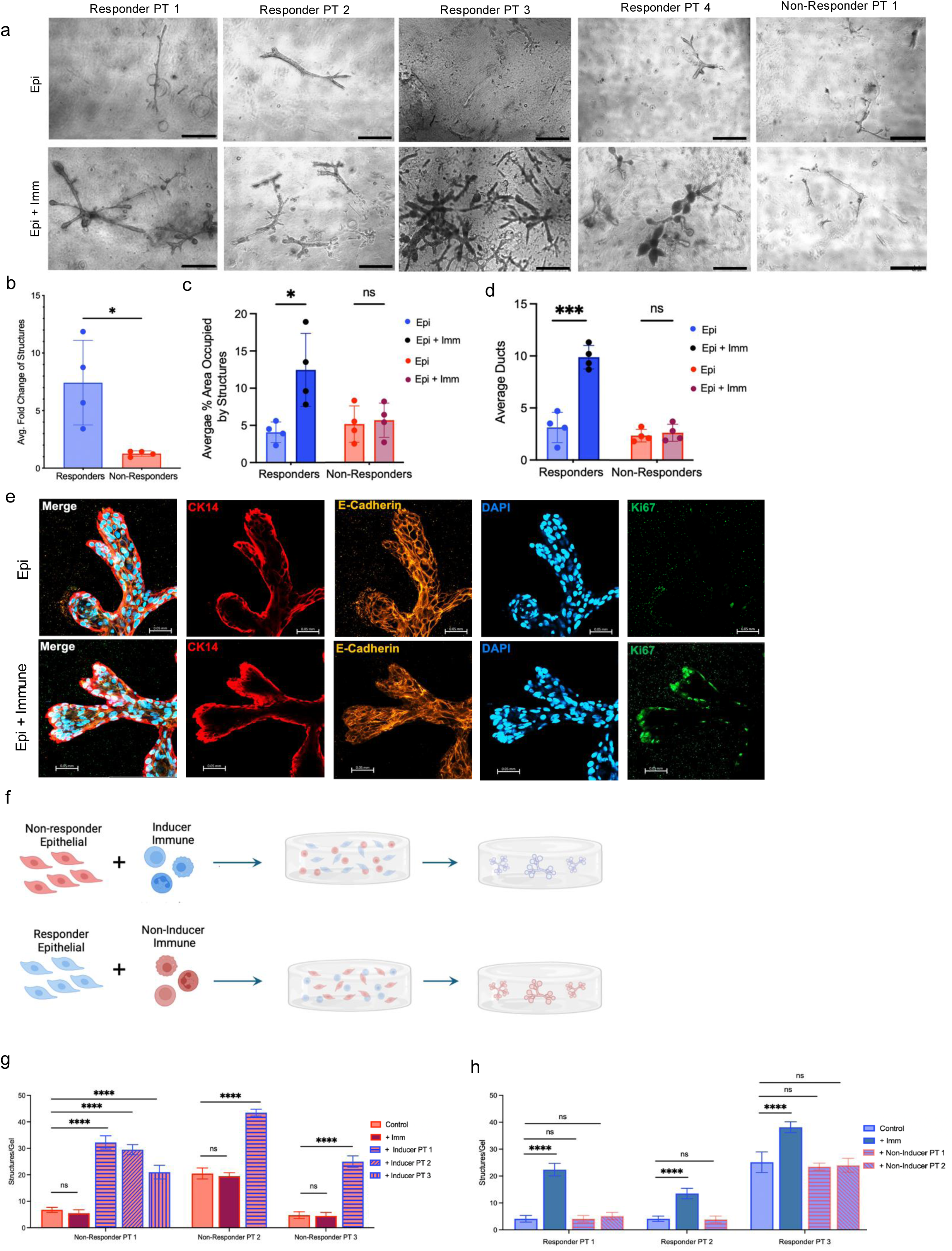
Immune cells promote epithelial proliferation and ductal morphogenesis in a subset of patient samples. (a) Representative brightfield images of organoids from responder patient samples (n=4) and non-responder patient sample (n=1) cultured with and without immune cells. (b) Bar graph showing the average fold change of structures in the presence of immune cells across 8 hydrogels for responders (n=4) and non-responders (n=4). *p-value < 0.05, ns, not significant, two-tailed t-test. (c) Bar graph showing average percent area occupied by structures across 8 hydrogels with and without immune cells for responders (n=4) and non-responders (n=4). *p-value < 0.05, ns, not significant, multiple t-tests. (d) Bar graph showing average number of ducts across 8 hydrogels with and without immune cells for responders (n=4) and non-responders (n=4). ***p-value < 0.001, ns, not significant, multiple t-tests. (e) Immunofluorescent (IF) images of co-cultured 3D hydrogels after 18 days of culture, stained with CK14 (red), E-cadherin (orange), DAPI (blue) for nuclei, and Ki67 (green). Scale bar = 500µm. (f) Experimental schematic for cross-patient epithelial and immune cell pairing. Non-responder epithelium was co-cultured with inducer immune cells, and responder epithelium was co-cultured with non-inducer immune cells. (g-h) Bar graphs showing that inducer immune cells convert non-responder epithelium to a responder phenotype, whereas non-inducer immune cells fail to do so across 8 hydrogels per sample (n=3 patients). ns, not significant, *p-value < 0.05, **p-value < 0.01, ***p-value < 0.001, ****p-value < 0.0001.

In 4 out of 8 donor pairs, the presence of immune cells significantly enhanced epithelial growth, with an average 4.7-fold increase in the number of organoids, with structures appearing at earlier time points, and the resulting organoids were both larger and more highly branched compared to those that formed by epithelial cells alone (Fig. 2a-d, Supplemental Fig. 3b). Ki-67 staining at day 18 confirmed that co-cultures maintained a significantly higher proliferative index compared to epithelial only cultures indicating that immune cells not only stimulate organoid formation but also sustain a heightened proliferative response even at late developmental timepoints (Fig. 2e). Interestingly, this effect was not universal to all donor pairs. In the remaining samples, immune cells failed to augment morphogenesis, prompting us to classify epithelial cultures as ‘responders’ or ‘non-responders’ and their cognate immune populations as ‘inducers’ and ‘non-inducers’, respectively (Fig 2a-d).

To determine whether the observed growth-promoting effect of paired donor samples was due to differences in the donor specific epithelial cells or in the immune cells, we co-cultured non-responder epithelial cells with inducer immune cells, and conversely, responder epithelial cells were co-cultured with non-inducer immune cells (Fig 2f). Reciprocal mixing experiments demonstrated that the growth-promoting capacity tracked with the immune compartment; inducer immune cells rescued non-responder epithelia, whereas non-inducer immune cells could not stimulate responder epithelial cells (Fig. 2f-h). These findings suggest that the functional heterogeneity among donor-derived immune cells, not intrinsic differences in epithelial cells, drives the variability in organoid growth. Furthermore, these results imply that immune cell composition, or secretory cytokine profiles are key regulators driving this effect.

### Cytokine Profile and Immune Cell Composition Underlie Differential Induction of Organoid Formation

Since inducer immune cells significantly enhanced organoid formation, while non-inducer immune cells had no effect, we reasoned that immune-cell-intrinsic features such as secreted cytokines, might drive the phenotype. Accordingly, we first compared the cytokine milieu generated by inducer versus non-inducer co-cultures. Supernatants from day 14 co-cultures were probed with an 80-plex human cytokine array. Inducer co-cultured secreted markedly higher levels of key cytokines, including CXCL5, GRO-α, LIF, IGFBP-2, MCP-1, IL-6, IL-8, and TIMP-1 (Fig. 3a). These cytokines are known to influence immune cell recruitment, epithelial proliferation, and tissue remodeling^18–29,29–35^. For example, MCP-1, CXCL5, and GRO-α are chemokines that facilitate immune cell chemotaxis, potentially enhancing the recruitment and activation of immune cells within the co-culture, thereby supporting epithelial cell proliferation^20,21,23,24,29,30,36^. LIF promotes cell survival and differentiation, playing a role in epithelial integrity and tissue remodeling^25,26^. IGFBP-2 regulates insulin-like growth factor availability, which is critical for epithelial growth^27,28^. IL-6 and IL-8 are pro-inflammatory cytokines that promote immune cell activation and epithelial proliferation^18,22,33,35^, while TIMP-1 regulates extracellular matrix remodeling and tissue repair^31,32^. In contrast, non-inducer co-cultures showed lower levels of these factors.

**Figure 3:**
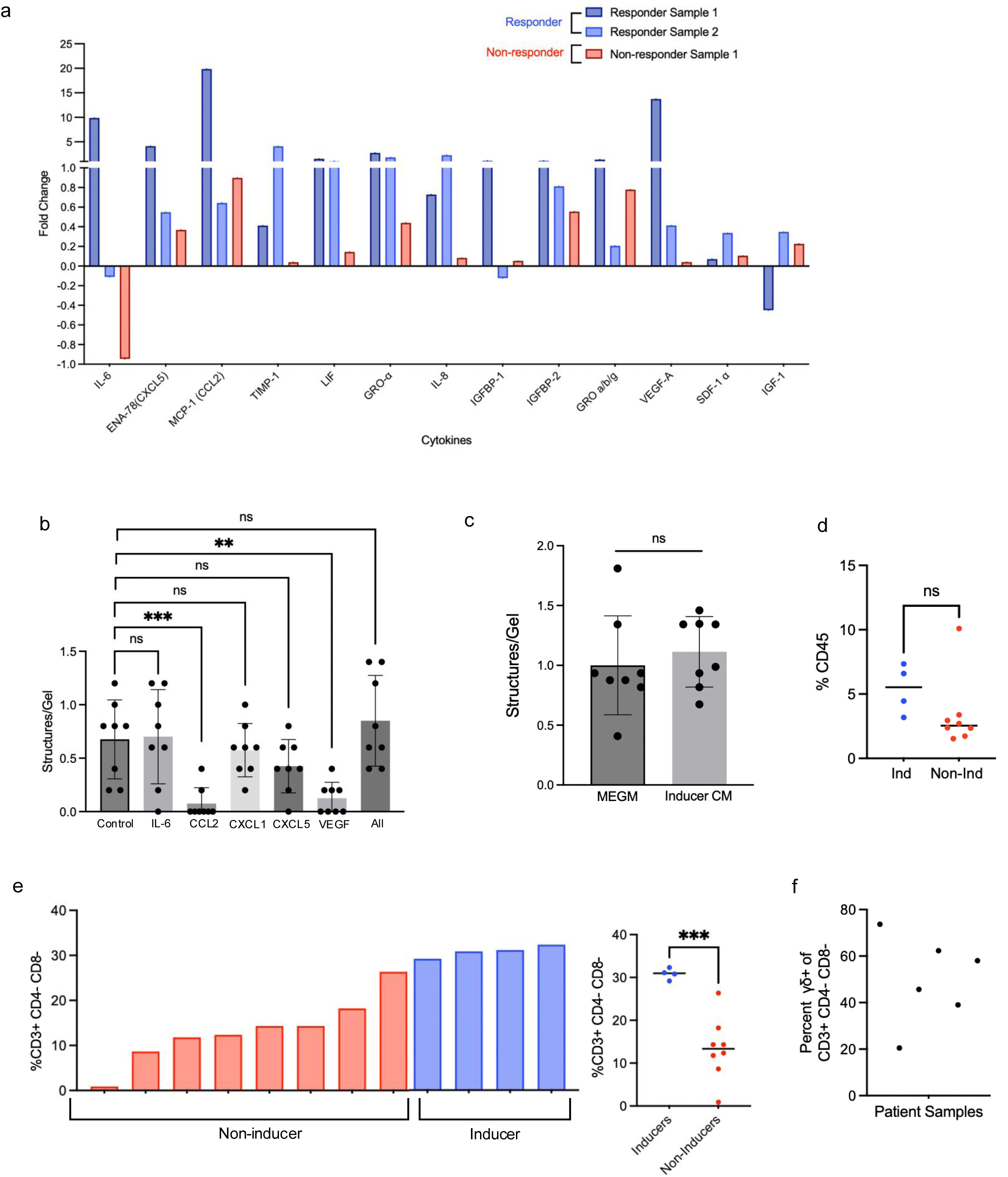
Immune cell presence is required to drive the epithelial growth phenotype, independent of soluble cytokine signals. (a) Cytokine array profiling comparing co-cultures and epithelial-alone hydrogels shows elevated levels of IL-6, CXCL5, CCL2, TIMP-1, LIF, GRO-a (CXCL1), IL-8, IGFBP-1, IGFBP-2, VEGF-A, SDF-1a, and IGF-1in responder co-cultures with immune cells (n=2 patients). (b) Bar graph showing treatment of responder epithelial cells with individual (IL-6, CCL2, GRO-a (CXCL1), CXCL5, VEGF) or pooled recombinant cytokines at 2ng/ml. (n=8; ns, not significant, **p-value < 0.01, ***p-value < 0.001). (c) Bar graph showing treatment of responder epithelium with conditioned media from responder co-cultures (n=8, ns, not significant). (d) Flow cytometry analysis of patient single cells showing percent of CD45 immune cells out of all breast cells. (n=12; ns, not significant). (e) Bar graph and scatter plot showing percent of CD3^+^ CD4^-^ CD8^-^ double negative (DN) T cells out of all CD45^+^ immune cells (n=12; ***p-value < 0.001, two-tailed t-test). (f) Scatter plot showing percent of γδ T cells out of all DN T cells.

To determine whether these soluble factors were causative, we treated non-responder epithelial cultures with the individual recombinant cytokines with individual recombinant cytokines CXCL5, GRO-α, VEGF, IL-6, or MCP-1. None of the individual cytokines alone could replicate the growth-promoting effects observed with inducer immune cells (Fig 3b). In addition, combining all the cytokines together also failed to reproduce the inducer immune cell phenotypes (Fig. 3b). Finally, conditioned media from inducer co-cultures added to non-responder cultures also failed to enhance organoid outgrowth as seen with inducer cells. Therefore, while inducer cultures create a pro-morphogenic cytokine environment, soluble factors alone do not account for the full inductive effect (Fig. 3c).

Given these results, we next profiled the cellular composition of tissue-resident immune populations by flow cytometry (n-12). CD45^+^ cells constituted 1-10% of total breast-derived cells, with significant patient heterogeneity observed across donors (Fig. 3d, Supplemental Fig. 3a). We observed no significant differences in the percentage of CD45+ cells between inducer and non-inducer samples. However, the inducer cohort contained a markedly larger fraction of double negative T cells (30.9% vs 11.5%; Fig. 3e). Within this compartment, γδ T cells represented 25-75% of the CD3^+^ CD4^-^ CD8^-^ cells (Fig. 3g, Supplemental Fig. 3c). Given the roles of γδ T cells in epithelial remodeling and immune surveillance, their enrichment suggests that the enhanced stimulation observed by inducer cells may be due to the presence of γδ T cells.

### Live-Cell Imaging Reveals Dynamic Immune Cell Behavior During Organoid Induction and Niche Formation

Given that soluble factors alone did not recapitulate the inducer phenotype, we hypothesized that direct physical interactions and dynamic cellular behaviors between responder immune cells must be critical. The single cell-derived 3D hydrogel system is uniquely suited to visualize such events, because organoid formation begins with dispersed stem-like epithelial cells that must build their own niche and elaborate TDLU architecture. This occurs though a well described stepwise process of induction and patterning, where single stem cells begin to differentiate and form the basic structures of the breast tissue^37–40^. By coupling the co-culture model with live imaging, we can observe in real-time the influence of inducer immune cells on the initial niche formation and differentiation of stem cells during the early days of culture. We can also visualize the rapid growth phases of morphogenesis and maturation when stem cell progeny undergo significant expansion including the formation of terminal end buds and their invasion during ductal elongation.

To capture these early inductive events, we dual labeled donor-matched immune and epithelial cells (Cytopainter green for CD45^+^ cells, Cytopainter deep red for epithelial cells) and performed confocal time-lapse microscopy for the first 7 days after seeding, which is the window when gross organoid growth is minimal, but lineage commitment (induction) and patterning decisions are initiated. At timepoint 0, the two cell populations (green-stained CD45^+^ and red-labeled epithelial cells) were evenly interspersed. By 24 hours, immune cells had begun directional migration, making transient contact with epithelial cells throughout the hydrogel. By day 7 immune cells had localized around the epithelial cells (Fig. 4a-c). Motility peaked between 24-48 hours and then progressively decreased as immune cells settled into stable positions abutting epithelial clusters (Fig. 4b, Videos 1, 2). By day 7, immune cells had formed a contiguous ‘halo’ around each nascent organoid, marking the first evidence of a self-assembled organoid (Fig. 4a, Supplemental Fig. 2a-c). The formation of supportive niches remained throughout the 28-day culture period, suggesting they may have a role beyond initial induction to influence morphogenesis and tissue maturation.

**Figure 4:**
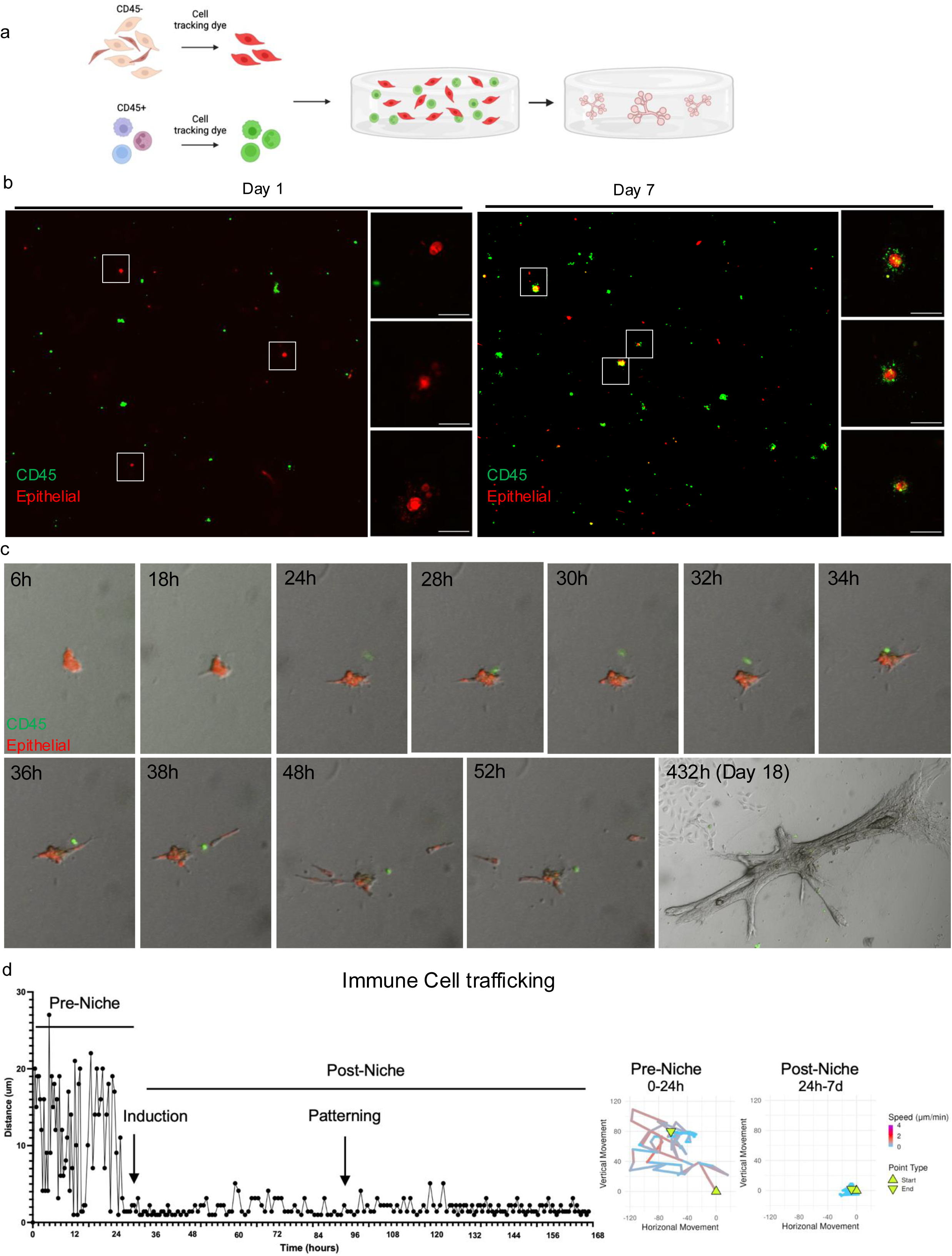
Immune cells dynamically localize to epithelial clusters, coinciding with early proliferation and morphogenesis. (a) Schematic of fluorescent labeling of epithelial cells (red) and immune cells (green) prior to hydrogel embedding. (b) Representative confocal images showing distribution of epithelial and immune cells at day 1 and clustering of immune cells adjacent to epithelial structures by day 7. Scale bars = 50 µm. (c) Spanel of selected frames from Supplemental Video 1, demonstrating dynamic interaction between immune and epithelial cells, with immune cells contacting epithelial clusters followed by epithelial cell division. (d) Quantification of motion and trajectory plots of immune cell for 0-24 hours and 24 hours-7 days of growth. The color gradient indicates the speed of the cell, and the triangles denote the starting and end points of the tracks. Quantification of immune cell motility demonstrates peak speed and displacement during the first 24–48 hours of culture.

Cultures containing non-inducer immune cells showed comparable motility but failed to establish this tight juxtaposition with epithelial cells (Videos 3-5). Rather, immune cells remained randomly distributed and disengaged from epithelial clusters throughout the imaging period. The absence of stable contacts preceded the failure of these cultures to accelerate morphogenesis, reinforcing the idea that timely immune-epithelial docking, and not simply cell trafficking, is required for induction. Taken together, the live-imaging data reveal a brief but important window in which spaciotemporal immune cues can sculpt breast tissue development. Early, contact-dependent interaction with immune cells appears to promote the subsequent surge in proliferation and the rapid development of ducts and lobules.

### TGF-β is a mediator of epithelial-immune cell-cell communication

The live imaging results revealed dynamic migration of immune cells towards epithelial cells during the early stages of organoid development, stimulation of induction, and evidence of immune cell localization and persistent niche formation. We therefore sought to identify the specific cell-cell communication mechanisms responsible for influencing epithelial cell fate decisions during this process. To achieve this, we leveraged two previously published single-cell RNA sequencing (scRNA-seq) datasets from both healthy human breast tissue^41^ and breast-tissue derived organoids grown in 3D hydrogels^42^. These datasets were integrated using Seurat v5^43^ to establish an overlap between in vivo cell populations and those observed in our 3D culture model (Fig. 5a).

**Figure 5:**
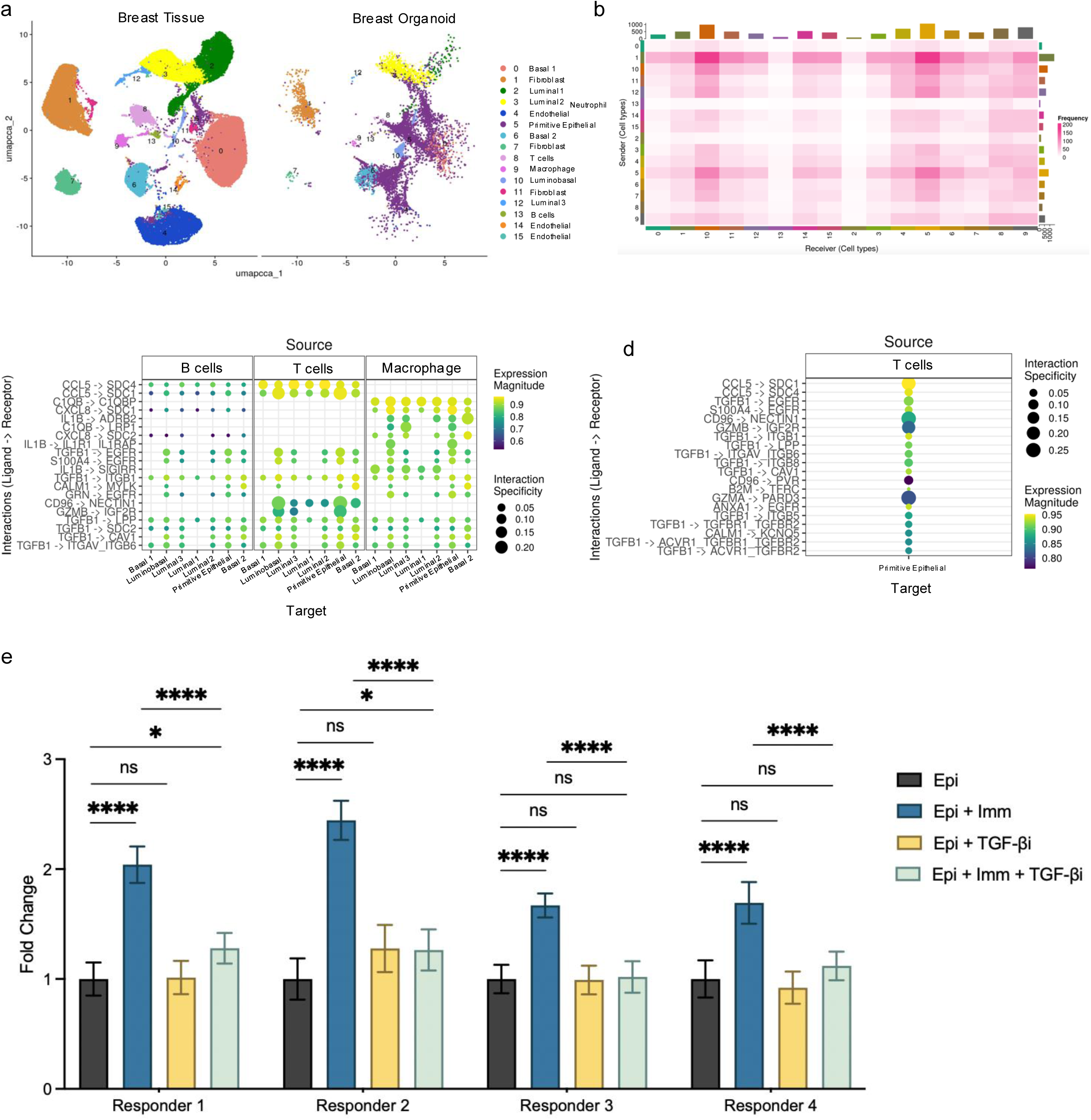
TGF-β signaling mediates immune-induced epithelial growth in responder samples. (a) UMAP projection of integrated scRNA-seq data from primary human breast tissue and hydrogel cultures. (b) Heatmap of inferred ligand–receptor interactions between all clusters. (c) Analysis of top 20 ligand–receptor pairs between immune cell clusters and epithelial cell clusters. (d) Analysis of the top 20 ligand-receptor pairs between T cells and primitive epithelial cells, showing 50% involve TGF-β signaling components. (e) Bar graph showing fold change of organoid number across responder sample conditions: epithelium alone, epithelium + immune cells, epithelium + TGF-β neutralizing antibody, and epithelium + immune cells + TGF-β neutralizing antibody (n=4 patients, 8 hydrogels each; ns, not significant, *p-value < 0.05, ****p-value < 0.0001).

Using this integrated dataset, we applied LIANA^44^ to infer ligand-receptor (L-R) crosstalk, focusing on immune-epithelial communications. Notably, we found that the primitive epithelial (Cluster 5) and Luminobasal epithelial population (Cluster 10) are communication ‘hot spots’ with some of the highest aggregate L-R scores with surrounding cell types (Fig. 5b). Among immune subsets, T cell (Cluster 8) and macrophages (Cluster 9) exhibited the strongest inferred interactions with the primitive epithelial (Cluster 5) (Fig. 5c). Strikingly, 10 of the top 20 T cell-primitive epithelial cell interactions involved the TGFB1 ligand (Fig. 5d). While the specific receptors varied (TGFBR1/2/3, ACVR1B), this recurring presence of TGF-β signaling suggested a prime candidate regulator for early fate decisions.

To directly test this, we treated epithelial cells with and without immune cells in the presence of a pan TGF-β neutralizing antibody or an IgG rabbit control. After 14 days of treatment, TGF-β blockade had no appreciable effect on responder cell organoid number or size (Fig. 5e). However, in the presence of inducer immune cells, TGF-β neutralization attenuated organoid formation, reducing the number of structures to levels comparable to epithelial-only cultures (Fig. 5e). These findings indicate that TGF-β plays a critical role in mediating the crosstalk between immune cells and epithelial cells during organoid development, and its activity is central to the immune niche formation.

## Discussion

In this study, we uncovered a previously underappreciated role for immune cells in driving breast tissue organogenesis via (i) rapid immune-epithelial docking that creates a physical niche, and (ii) T cell-derived TGF-β signals within that niche that trigger stem-cell activation and accelerated tissue assembly. By establishing a 3D hydrogel-based co-culture system, we were able to recapitulate key aspects of the human breast tissue microenvironment, allowing for real-time visualization of immune-epithelial interactions in human development that have been difficult to study in vivo. In addition, by mapping temporal and spatial dynamics through longitudinal studies and live-cell imaging, we were able to provide critical insights into how immune-epithelial crosstalk drives organoid development and regeneration.

Multiple subsets of T cells have been implicated in playing roles in shaping the epithelial niche in tissue homeostasis and repair^6–10,45–51^. Here, we show that immune cells, particularly those enriched in γδ T cells, could significantly accelerate breast organoid formation and maturation. Co-culture of epithelial cells enriched with this subset of T cells led to a marked increase in both the number and size of organoids compared to cultures without immune cells or with non-inducer immune cells. The enrichment of γδ T cells is particularly interesting, as these cells have unique properties that bridge innate and adaptive immunity and are increasingly recognized for their roles beyond classical immune functions, including tissue homeostasis, wound repair, and even developmental processes. In humans and rodents, γδ T cells are prevalent in the female reproductive tracts and fluctuate in response to hormone changes across the reproductive cycle^52^. This suggests these cells might be responding to and possibly reinforcing niche signals that drive stem-cell activation and lineage decisions that take place during each menstrual cycle. This observation, combined with our results, suggests that the immune compartment not only stimulates epithelial proliferation, as evidenced by increased Ki-67 expression, but also plays a crucial role in orchestrating the spatial organization and niche formation that underlies proper tissue architecture. Elucidating the specific roles of these subsets requires targeted approaches, such as single-cell transcriptomics, functional assays, and depletion studies.

While cytokine profiling experiments indicate that the enhanced organoid formation is associated with elevated secretion of several cytokines known to influence tissue remodeling and proliferation, including IL-6, IL-8, and MCP-1, cytokines alone were not able to mimic the immune-induced phenotype. Rather, live imaging showed direct cell-cell interactions, and the physical presence of immune cells, was critical for initiating and sustaining the complex signaling networks that drive organogenesis. This was observed within the first 24 hours in the dynamic behavior of immune cells during the early stages of organoid induction followed by their stable localization in emerging epithelial structures, highlighting their role as active participants in niche formation.

The dynamic interplay between immune cells and epithelial cells, was mediated in part by TGF-β signaling. Inhibition of TGF-β signaling selectively abrogated the growth-promoting effects of inducer immune cells without impacting epithelial growth in monocultures. This suggests that TGF-β functions as a key effector in the paracrine and possibly contact-dependent interactions that facilitate both the induction of epithelial proliferation and the formation of a supportive niche.

While TGF-β is a well characterized cytokine involved in immune regulation, inflammation, and tissue remodeling, its role in immune-mediated epithelial induction during organogenesis remains underexplored, particularly in the context of human breast tissue development^53–56^. Here, we found that TGF-β from immune cells was required to directly stimulate the induction and proliferation of epithelial cells in a controlled organoid model. This suggests that immune derived TGF-β is not just involved in later stages of mammary development, tissue maintenance, or repair but is also crucial for the early induction phases of development. TGF-β is also a pleiotropic cytokine with both proliferative and inhibitory roles, depending on the context^54–59^. It is possible that TGF-β from immune cells in this context modulates epithelial stem cell states, priming cells for expansion and differentiation through downstream pathways such as SMAD signaling, Wnt activation, or Hippo pathway modulation^53,60–62^.

Future studies aimed to identify the specific immune cell subsets and molecular signals driving this reprogramming will be important to better understand which immune components selectively enhance epithelial growth. Moreover, defining the precise epithelial stem and progenitor cell populations that respond to immune cues will provide insight into how immune cells shape the epithelial hierarchy and influence developmental trajectories. In addition, many of the same immune-epithelial interactions that enhance normal tissue growth may be co-opted in the tumor microenvironment, where immune cells can either promote or suppress tumor progression, with TGF-β serving as a prime example. It is well-documented as a context-dependent mediator of tumorigenesis, acting as both a suppressor in early stages and a promoter in advanced cancers^63–65^. Understanding how TGF-β signaling operates within the immune niche during normal development could provide key insights into its dysregulation in cancer.

### Conclusion

Our study highlights the immune niche as a central regulator of epithelial stem cell dynamics, expanding our understanding of how immune cells shape tissue development and homeostasis. The heterogeneity in responder and non-responder phenotypes underscores the importance of defining patient-specific immune signatures and understanding their impact on epithelial cell behavior. Future research should prioritize identifying the immune subsets and signaling pathways responsible for promoting epithelial growth, with a focus on TGF-β signaling, direct cell-cell interactions, and their convergence in modulating epithelial cell fates. By bridging these insights, we have the potential to inform regenerative medicine approaches and uncover novel therapeutic strategies for breast tissue repair and disease treatment.

## Methods

### Ethics Statement

Primary tissues that normally would have been discarded as medical waste post-surgery were obtained in compliance with all relevant laws, using protocols approved by the institutional review board at Maine Medical Center and Tufts Medical Center. All tissues were anonymized before transfer to prevent tracing back to specific patients; for this reason, this research was provided exemption status by the Committee on the Use of Humans as Experimental Subjects at the Massachusetts Institute of Technology, and at Tufts University Health Sciences (IRB# 13521). All enrolled patients in this study signed an informed consent form agreeing to participate in this study and for publication of the results.

### Tissue processing and 3D culture

Tissues were partially dissociated with 1.5mg/ml collagenase (10103578001, Millipore Sigma) and 18μg/ml hyaluronidase (H3506, Sigma Aldrich), yielding small epithelial fragments, washed, and cryopreserved in 1 mL aliquots. Tissue fragments were further dissociated to single cells using 0.25% trypsin (25200056, Thermo Fisher Scientific) and 5 U/ml Dispase II (4942078001, Roche). Immune cells were isolated using the EasySep release human CD45 positive selection kit (100-0105 Stem Cell Technologies). Hydrogels were seeded with epithelial cells at a concentration of 1000 cells per 100 μL gel, and immune cells were seeded at a concentration between 6,500-10,000 cells per 100ul gel.

For staining of cells for live imaging, cells were stained at a 1:500 concentration of Cytopainter green (ab138891, Abcam) or Cytopainter deep red (ab138894, Abcam) in PBS for 30 minutes at 37 °C. Cells were washed 2x in PBS and 1x in MEGM prior to seeding.

For hydrogel fabrication and seeding, cells were resuspended in 1.7 mg/ml collagen I (08-115, Millipore Sigma), 10 μg/ml hyaluronic acid (385908, Millipore Sigma) 40 μg/ml laminin (23017-015, Gibco), and 20 μg/ml fibronectin (F1056, Sigma Aldrich), pH 7.3. Gels were deposited in 8-well chamber slides (354118, Falcon) and incubated 1 hour at 37 °C for polymerization before adding MEGM medium. Hydrogels were cultured floating in serum-free mammary epithelial growth media (MEGM) in 8-well chamber slides. MEGM consisted of basal medium (M171500, Thermo Fisher Scientific) supplemented with 52μg/ml bovine pituitary extract (13028014, Thermo Fisher Scientific), 10ng/ml hEGF (E9644, Sigma Aldrich), 5μg/ml insulin (I9278, Sigma Aldrich), 500ng/ml hydrocortisone (H0888, Sigma Aldrich), 1% v/v GlutaMAX (35050061, Thermo Fisher Scientific), and 1% v/v penicillin-streptomycin antibiotics (15140122, Thermo Fisher Scientific).

For treatment with TGF-β neutralizing antibody, hydrogels were cultured with the antibody (AB-100-NA, R&D Systems, 2ug/ml) in MEGM from day 0-14, and then released for the remainder of the culture period.

For treatment with conditioned media, conditioned media was harvested from hydrogel cultures, filtered through a 0.2um filter, and snap frozen and stored at −80°C. Conditioned media was mixed at a 1:1 ratio with fresh MEGM media and added to cultures at timepoint 0 and continued for the entire culture period.

For treatment with recombinant proteins IL-6 (PHC0066, Thermofisher,2ng/ml), CCL2 (279-MC-050/CF, R&D Systems, 2ng/ml), GRO-a (275-GR-010/CF, R&D Systems, 2ng/ml), CXCL5 (254-XB-025/CF, R&D Systems, 2ng/ml), VEGF (78073, Stem Cell Technologies, 2ng/ml), treatment began at timepoint 0 and continued for the entire culture period.

### Live Imaging

Primary single cells were incubated with the cell tracking dye as described above, and seeded at a concentration of 100 cells per 20ul hydrogel. 20ul hydrogel drops were deposited into the center wells of a 96-well plate (3603, Corning). Hydrogels were incubated at 37 °C for 1 hour until polymerized. 80ul of MEGM media was added to each well and the hydrogels were lifted off of the well surface with a pipette tip. Cultures were then placed into a pre-warmed incubation chamber (Okolab inc) enclosed over a Nikon Eclipse Ti2-AX confocal microscope (Nikon Microscopy). Images of selected points of whole hydrogels were collected starting immediately after the addition of media, and every 30-45 minutes after, in brightfield and with AF488 and AF647 lasers at 4x magnification and 1.6x zoom across 5-9 z-positions. 20-40ul of MEGM media was added to the culture twice a week to maintain proper growth factors and liquid volume to prevent hydrogels from drying out. Cultures were live imaged for 7-21 days. Analysis and production of videos across locations and timepoints was performed using NIS-Elements (Nikon) and iMovie (Apple) software.

### Individual Cell Tracking

Using FIJI software, individual cells were tracked across videos of live confocal imaging, depicting the development of organoids from single cells. The x/y coordinates of cells were manually annotated frame-by-frame to generate precise tracking data. The speed of cells were calculated for each segment by measuring the displacement over the time interval between successive frames. The frame rate of the video was 35-55 minutes (precise times were known for each frame) allowing high temporal resolution. Visualization of cell tracks and speed measurements was done using R (Version 4.3.3) utilizing dplyr and ggplot2 packages. Adjustments were made to cell coordinates by anchoring the starting points at (0, 0) to show clear visualization of movement paths. The positioning of cells was determined in each frame, and the speed was deduced by the elapsed time between frames.

### Immunofluorescence Staining

Hydrogels were fixed with 4% paraformaldehyde (15710-S, Electron Microscopy Sciences) at room temperature for 30 minutes and washed in PBST, comprised of 1X PBS (46-013-CM, Corning) and 0.05% Tween 20 (P7949, Millipore Sigma). Samples were then permeabilized in 0.1% TritonX-100 (X100, Sigma Aldrich) in PBST and incubated with blocking buffer comprised of PBST with 3% BSA (BSA-50, Rockland Immunochemicals) and 10% goat serum (G9023, Millipore Sigma) for 2 hours at room temperature. Hydrogels were then stained with the appropriate primary antibody in blocking buffer at 4°C overnight. Samples were washed with PBST and incubated with secondary antibody for 2 hours at room temperature, and DAPI for 30 minutes at room temperature. Primary antibodies used in this study were: E-cadherin (ab1416, Abcam, Clone HECD-1, 1:100), CK14 (RB-9020, Thermo Fisher, 1:300), CD45 (04-1102, Milipore, 1:500), CD3 (344802, Biolegend, 1:500), Ki-67 (ab254123, Abcam, 1:500). Secondary antibodies used in this study were: goat-anti-mouse-AF555 (A21424, Life Technologies, 1:1000), goat-anti-rabbit-AF488 (A11008, Life Technologies, 1:1000), goat-anti-rabbit-AF647 (A21245, Life Technologies, 1:1000), goat-anti-chicken-AF488 (A11039, Life Technologies, 1:1000). Dyes and probes used for immunofluorescence: Phalloidin-AF647 (A22287, Life Technologies, 1:250 of 400x stock), DAPI (D1306, Life Technologies, 5μg/ml).

### FFPE Patient Tissue Staining

Patient tissue was fixed in 10% buffered formalin and paraffin-embedded in tissue blocks which were serially sectioned. Slides were baked at 55°C for 45 min, then rehydrated by sequential incubation in Xylene, 100% ethanol, 95% ethanol, 70% ethanol, and 50% ethanol for 3 minutes each, then 10 minutes in PBS. Antigen retrieval was done in sodium citrate buffer in a steamer for 20 minutes. Slides were washed in TBS-T comprised of 1X TBS (Trizma base, T4661, Millipore Sigma; NaCl, 71376, Millipore Sigma) and 0.25% TritonX-100 and then in blocking buffer, comprised of TBS-T with 10% goat serum and 1% BSA, for 2 hours at room temperature. Endogenous avidin and biotin were blocked for 15 minutes and washed with TBS-T. Slides were stained with primary mouse-anti-human CD45 antibody (14-0459-82, clone HI30, eBioscience, 1:150) in blocking buffer at 4°C overnight. Samples were washed with TBS-T and incubated with secondary biotinylated goat-anti-mouse antibody (BA-9200, Vector, 1:200) for 2 hours at room temperature. Vectastain ABC solution (Vector, PK-4000) was prepared and incubated for 30 minutes during wash steps. ABC was added to slides and incubated for 1 hour at room temperature. Slides were washed with TBS-T and 1x DAB was added to slides for 10 minutes. Slides were washed with TBS-T and hematoxylin was added for 5-10 minutes. Slides were washed in tap water 0.1% bicarbonate. Slides were dehydrated by sequential incubation in 95% ethanol, 100% ethanol, and Xylene. Permount (SP15-100, Fisher) mounting medium was added and covered with glass coverslips.

### Microscopy

Immunofluorescence images were captured using a Nikon AXR (Nikon Microscopy) or a Zeiss LSM800 scanning confocal microscope (Zeiss Inc). Brightfield images were captured using a Nikon AXR (Nikon Microscopy) or a Nikon Eclipse Ti-U (Nikon Microscopy) using SPOT 5.6 software.

### Cytokine Array

Conditioned media was harvested from hydrogel cultures, filtered through a 0.2um filter, and snap frozen and stored at −80°C. The human C5 cytokine array membrane (AAH-CYT-5-8, RayBiotech) was blocked with 2ml Blocking Buffer for 30 minutes at room temperature on orbital rocker. Blocking Buffer was aspirated from membrane and 1ml of conditioned media was added and membranes were incubated at 4°C overnight on orbital rocker. Membranes were washed in 2ml of 1x Wash Buffer I three times, and then membranes were washed with 2ml of 1x Wash Buffer II two times. 1ml of the biotinylated antibody cocktail was added to the membranes and incubated for 1.5 hours at room temperature. Membranes were washed in 2ml of 1x Wash Buffer I three times, and then membranes were washed with 2ml of 1x Wash Buffer II two times. Membranes then received 2ml of 1x HRP-Streptavidin and incubated for 2 hours at room temperature and then washed stated above. 500ul of the detection buffer was added to membranes and then imaged with the Chemidoc XRS+ with Image Lab 6.0.1 software (BioRad, Hercules, CA). ImageJ2 (Version 2.8.0/1.53t) with the Protein Array Analyzer package used for quantifications.

### scRNA-seq analysis

We integrated and analyzed previously published scRNA-seq data from organoids grown in ECM hydrogels (CITE) and healthy primary breast tissue. The data was analyzed using Seurat V5 for data integration, normalization, and feature selection. Briefly, 10X files were loaded and Seurat objects were generated for each data set. Filtering removed cells with < 200 or ώ7500 genes and with mitochondrial content >7.5%. Genes detected in less than 3 cells were dropped from analysis. The Seurat objects were merged using the merge function, and the data was then normalized by multiplying transcripts by a factor of 10,000 and then log-transforming the data. Variable features used for analysis were identified by using the FindVariableFeatures function at default settings. Data was then scaled and PCA analysis was run. Integration was done with the CCAIntegration method using the IntegrateLayers function in Seurat. Cells were then clustered using K-nearest neighbor (KNN) graphs and the Louvain algorithm using the first 15 dimensions from the dimensional reduction analysis. Clusters were called using the FindClusters function with a resolution of 0.3, and 16 distinct cell clusters were identified. Clustered cells were visualized by UMAP embedding using the default settings in Seurat. The layers of the integrated Seurat object were then joined using the JoinLayers function. To identify differentially expressed genes between cell clusters, we utilized the FindAllMarkers function to identify features detected in >10% of a cell cluster compared to all other cells. Pathway analysis to identify enriched biological pathways associated with differentially expressed genes was done using established databases, such as PanglaoDB. The top differentially expressed markers were used to determine gene expression location and assign cluster names.

### Cell-cell inference

Cell-cell communication inference was done using the integrated scRNA-seq data and the LIANA R package. Briefly, LIANA was run using the liana_merge function using the integrated Seurat object. Results were aggregated using the RobustRankAggregate method using the liana_aggregate function. The top 20 interactions are unique ligand-receptors ordered consequentially.

### Flow Cytometry

Single cells dissociated from patient tissues as described above were washed with PBS with 2% FBS and stained with antibodies at 4°C for 60 minutes. Cells were then washed and resuspended in PBS with 2% FBS for flow cytometry. All data acquisition was done using an LSR II (BD Biosciences) and analyzed using FlowJo software (TreeStar). Antibodies used for flow cytometry: CD45-APC (555485, BD Biosciences), CD45-PE (555483, BD Biosciences), CD8-APC (566852, BD Biosciences), CD8-FITC (322704, BioLegend) CD3-APC-Cy7 (557832, BD Biosciences), CD4-PacBlue (344619, BD Biosciences), TCR γ/δ-FITC (331207, BD Biosciences), 8-color human immunophenotyping kit (130-120-640, Miltenyi).

### Statistics

All statistics were performed using GraphPad Prism 8-10. Student’s t-tests (two sided) were performed as a determinant of significance unless otherwise stated. Data expressed as Mean ± SD. Significance levels indicated as follows: * p-value < 0.05, ** p-value < 0.01, *** p-value < 0.001, **** p-value < 0.0001.

## Supplemental Figure Legends

**Supplemental Figure 1: Immune cells localize to developing epithelial structures in 3D hydrogel cultures.**

(a) Immunofluorescent (IF) images from representative patient-matched co-cultured 3D hydrogels after 21 days of culture, stained with DAPI (blue) for nuclei, CD45 (green) for immune cells. Scale bar = 500µm. (b) Immunofluorescent (IF) images from representative patient-matched co-cultured 3D hydrogels after 21 days of culture, stained with phalloidin (red) for actin cytoskeleton, DAPI (blue) for nuclei, CD45 (green) for immune cells, and CD3 (orange) for T cells. Scale bar = 500µm. (c) Immunofluorescent (IF) images from representative patient-matched co-cultured 3D hydrogels after 21 days of culture, stained with phalloidin (red) for actin cytoskeleton, DAPI (blue) for nuclei, CD45 (green) for immune cells, and CD3 (orange) for T cells. Scale bar = 50µm.

**Supplemental Figure 2: Validation of immune cell labeling and early localization to epithelial clusters in 3D co-culture.**

(a) Flow cytometry histogram and bar graph demonstrating strong signal and high viability of immune cells stained with Cytopainter green with minimal toxicity. (b) Gating strategy confirming robust and selective staining of immune (CD45+) populations with Cytopainter green. (c) Representative immunofluorescence snapshots from live imaging (0–48 h) showing green-labeled immune cells localizing around red-labeled epithelial clusters shortly after embedding. Images are representative of at least 2 independent experiments. Scale bars = 100μm.

**Supplemental Figure 3: Immune composition of patient samples and quantification of epithelial response by immune subtype.**

(a) Flow cytometry analysis showing the immunophenotype of CD45⁺ immune cells from patient breast tissue, including proportions of T cell, B cell, and myeloid subsets (n=12 patients). (b) Quantification of total epithelial structures per hydrogel, normalized to 1.0 for epithelial-alone controls, across 4 responder and 4 non-responder patients with and without immune cells. Data represent mean ± SEM. (c) Representative gating strategy for identifying γδ T cells within the CD3⁺CD4⁻CD8⁻ double-negative T cell population.

## Supplemental Video Legends

**Supplemental Video 1:** This time-lapse video is of a responder patient sample and shows a green-labeled immune cell directly interacting with a red-labeled epithelial cell cluster during the first 48 hours of co-culture within the hydrogel. The video then follows epithelial morphogenesis over an 18-day period. Framed were acquired every 30-45 minutes during growth.

**Supplemental Video 2:** This Time-lapse video is of a responder patient sample and shows a green-labeled immune cell localizing to and interacting with a red-labeled epithelial cluster in 3D hydrogel during the first 48 hours of co-culture. Playback is slowed to highlight dynamic cellular movements and early contact events. Frames were acquired every 30-45 minutes during growth. Same field of view as Supplemental Video 1

**Supplemental Video 3:** Time-lapse video of a non-responder sample showing red-labeled epithelial cells and green-labeled immune cells dispersed throughout the 3D hydrogel during the first 72 hours of growth. Small green immune cells exhibit rapid, erratic movement during the first 24 hours, with minimal motion observed during the final 24 hours of the 72-hour imaging period. Images acquired every 30–45 minutes.

**Supplemental Video 4:** Time-lapse video from a non-responder co-culture showing a red-labeled single epithelial cell expanding into an organoid over 21 days in 3D hydrogel. Green-labeled immune cells remain distant, with no evidence of direct contact or coordination. Images acquired every 30-45 minutes.

**Supplemental Video 5:** Time-lapse video showing red-labeled epithelial cell expansion and organoid formation over 21 days in a second non-responder patient sample. No direct interaction with green-labeled immune cells is observed throughout the imaging period. Images acquired every 30-45 minutes.

