## Supplementary figures and images for "Immune-Epithelial Interactions via TGF-β Orchestrates Stem-Cell Niche Formation and Morphogenesis"

### Supplmental Figures 1-3

Supplemental Figure 1

a

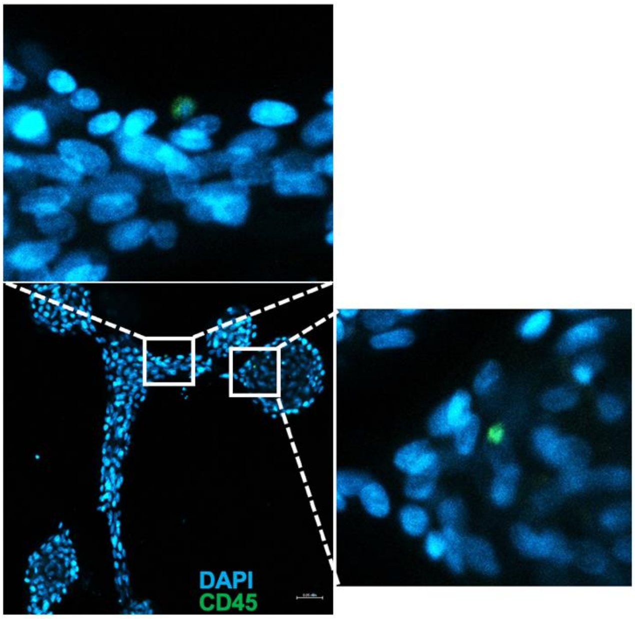


b

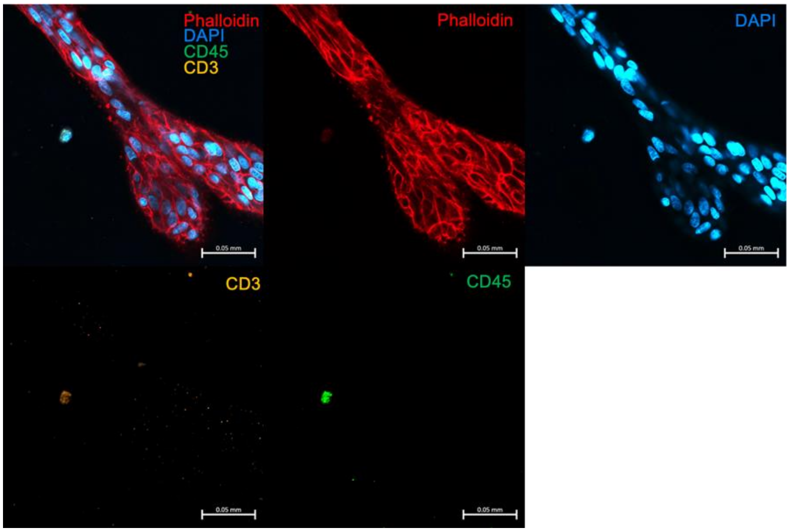


c

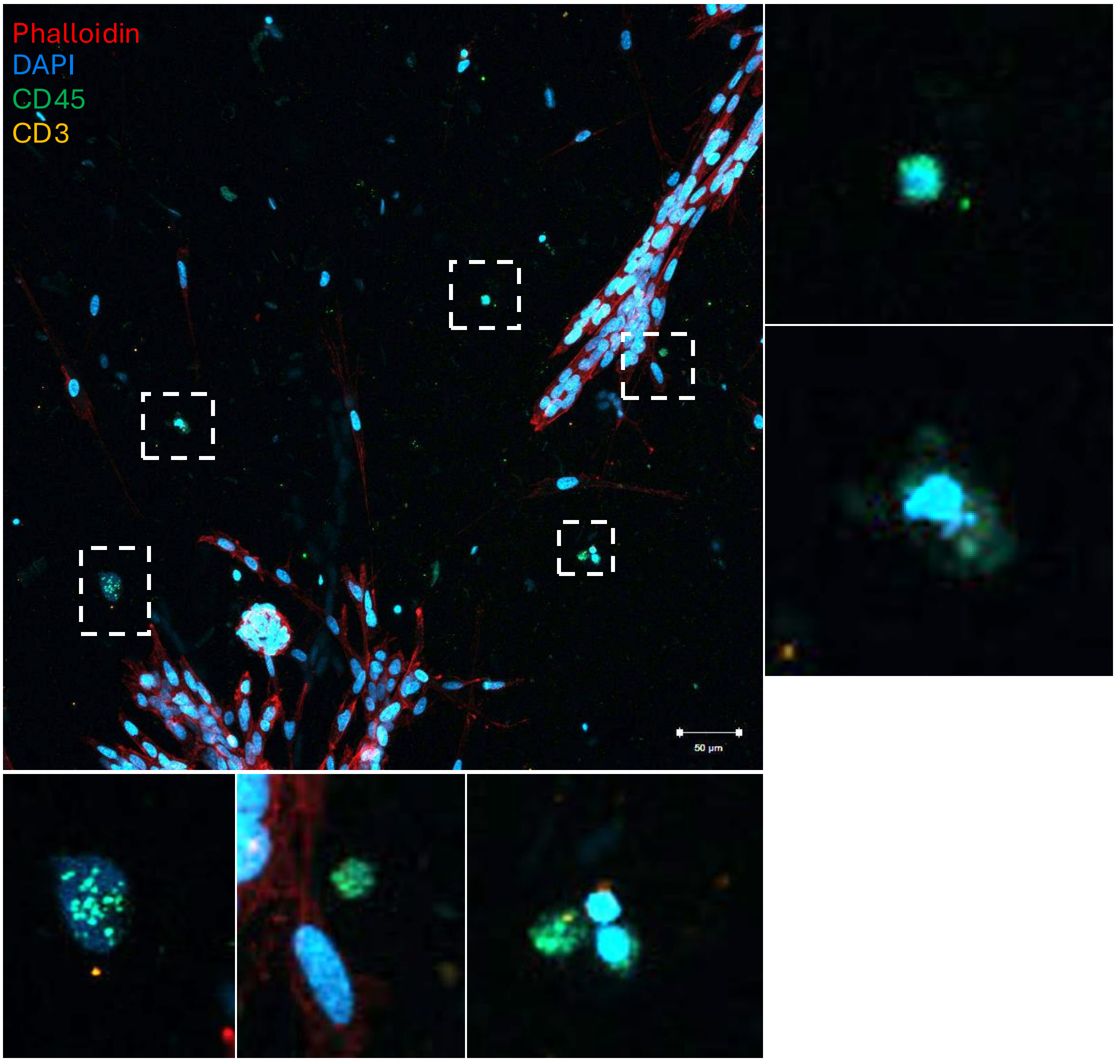

Supplemental Figure 2

a

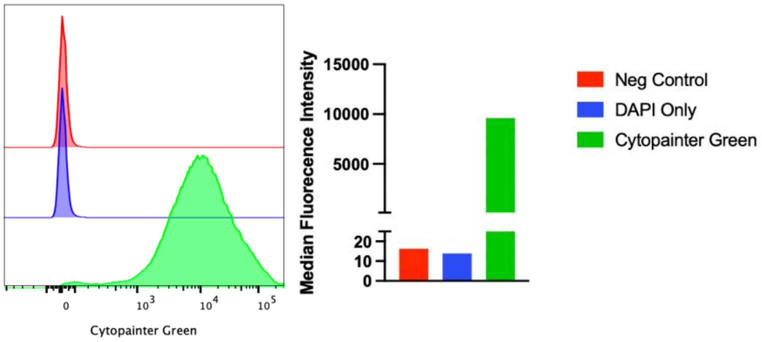


b

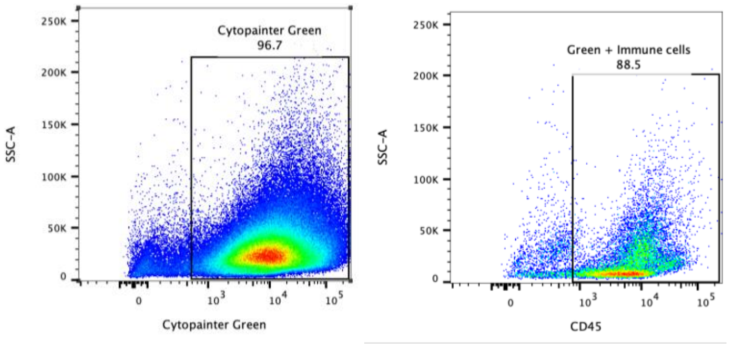


c

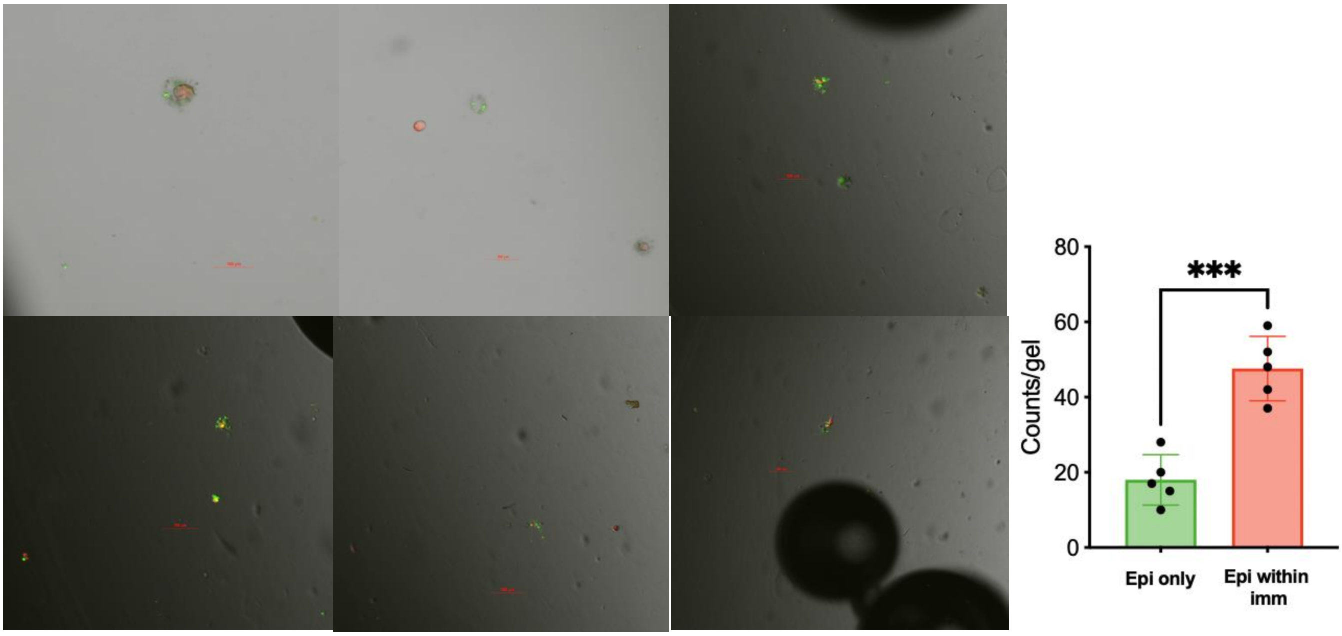


b

Supplemental Figure 3

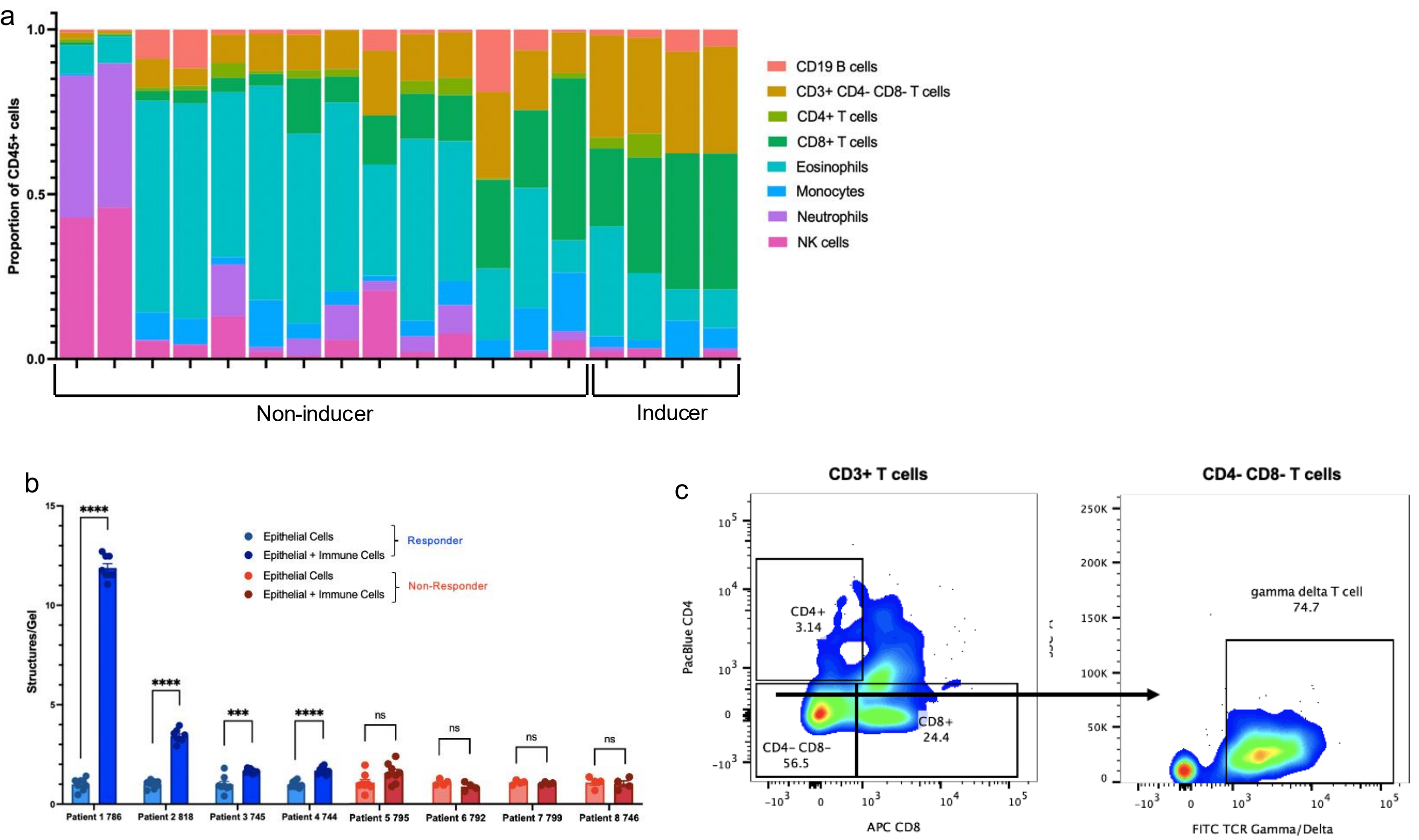
